# Tagging of Exo70 at the N-terminus compromises its assembly with the exocyst complex and changes its spatiotemporal behavior at the plasma membrane

**DOI:** 10.1101/2024.04.21.590474

**Authors:** Hiteshika Gosain, Guiscard Seebohm, Michael Holtmannspötter, Rainer Kurre, Karin B. Busch

**Author notes:** corresponding author Karin B. Busch.

## Abstract

The vesicle-tethering exocyst complex is a key regulator of cell polarity. The subunit Exo70 is required for the targeting of the exocyst complex to the plasma membrane. While the N-terminus of Exo70 is important for its regulation by GTPases, the C-terminus binds to PI(4,5)P2 and Arp2/3. Here, we compare N- and C-terminal tagged Exo70 with respect to subcellular localization, dynamics and function in cell membrane expansion. Using high-resolution imaging, we determined the spatial distribution and dynamics in different sub-compartments of un-polarized and polarized cells. With lattice light-sheet microscopy, we show that HaloTag-Exo70, but not Exo70-HaloTag, promotes the outgrowth of filopodia-like structures from the axon of hippocampal neurons. Fluorescence lifetime imaging of sfGFP-Exo70 and molecular modeling results suggest that the assembly of sfGFP-Exo70 with the exocyst complex is reduced. This is supported by single particle tracking data showing higher mobility of N- than C-terminal tagged Exo70 at the plasma membrane. The distinct spatiotemporal properties of N-terminal tagged Exo70 were correlated with pronounced filopodia formation in unpolarized cells and neurons, a process that is less reliant on exocyst complex formation. We therefore propose that N-terminal tagging of Exo70 shifts its activity to processes that are less exocyst-dependent.

**Why it matters:** In the life sciences, the high-resolution visualization of processes in living cells is of great interest. To tag proteins, they are fused with fluorescent proteins, usually at the N- or C-terminus of the amino acid sequence. We show here that the position of the tag alters the function and interaction of Exo70, a polypeptide involved in vesicle fusion, but also in membrane bending and expansion. Using state-of-the-art microscopic techniques, single particle localization and tracking, fluorescence lifetime imaging microscopy and co-localization in combination with modeling, we conclude that N-terminal tagging of Exo70 impairs its assembly with the exocyst complex and instead promotes the interaction of free Exo70 with the actin skeleton, which favors controlled membrane expansion into filipodia.

## 1 INTRODUCTION

The exocyst complex is an octameric complex consisting of Sec3, Sec5, Sec6, Sec8, Sec10, Sec15, Exo70 and Exo84 ^1^. Exo84, Exo70, Sec10 and Sec15 form one tetrameric subcomplex, Sec3, Sec5, Sec6 and Sec8 form the second tetramer. The holo-exocyst complex is required for multiple cellular tasks linked to exocytosis ^2^. However, specific subunits, such as Exo70, have been linked to specialized functions that link exocytosis to cell adhesion, migration and tumor invasion ^3^. Exo70, together with Sec3, mediates the association of the exocyst complex to the plasma membrane (PM), which is a critical step for vesicle tethering ^4,5^.

The subunits of the exocyst complex display a rod-like structure, and the core of the hollow exocyst complex is formed by the N-termini of all subunits except Sec10 ^6^. The C-termini point outwards. Via its C-terminus, Exo70 interacts with the plasma membrane lipid phosphatidylinositol 4,5-bisphosphate (PI(4,5)P_2_) ^7^. The N-terminus of Exo70 is responsible for the interaction with GTPases TC10 and Cdc42b ^8,9^. Chopping the N-terminus produced a dominant-negative Exo70 mutant ^9^, while fusion of a tag to the N-terminus was associated with formation of filopodia like structures ^10,11^, likely by interacting with the local actin meshwork via the Arp2/3 complex ^11^.

To better understand the effect of N-terminal tagging on Exo70 dynamics and thus function at the membrane, we compared the spatiotemporal behavior of N- and C-terminal tagged Exo70 by single molecule Tracking and Localization Microscopy ^12,13^. Additionally, we used fluorescence lifetime imaging microscopy, FLIM, to investigate the nano-environment and electrodynamic interactions. N-terminally tagged Exo70 displayed a higher mobility than C-terminally tagged Exo70, had a longer lifetime and promoted filopodia formation in unpolarized cells and neurons. Modeling suggests that N-terminally tagged Exo70 does not assemble into the exocyst complex. This explains the different modes of action.

## 2 MATERIALS AND METHODS

### Cell culture and cultivation

HeLa cells (purchased from DSMZ, # ACC 57) were chosen as a nonpolar model cell line. HeLa cultures were incubated in a T25 flask containing pre-warmed growth medium at 37°C and 5% CO_2_ throughout the experiments. The growth media consist of Dulbecco’s Modified Eagle Medium (DMEM), 10% Fetal Bovine Serum (FBS), 1% non-essential amino acids (NEA), 1% HEPES buffer solution, 1% L-Glutamine and 1% Penicillin as an antibiotic. Table 1 lists general material used and Table 1 culture media composition.

**Table 1:** Cell culture media composition

| Media | Composition |
| --- | --- |
| DMEM++ (growth medium) | 1x DMEM with Phenol red, 10% heat-inactivated FBS, 2 mM L-Glutamine, NEAA (1%), HEPES (1 µM) and 50 µg/mL Penicillin streptomycin |
| DMEM+++ (imaging medium) | 1x DMEM without Phenol red, 10% heat-inactivated FBS, 2 mM L-Glutamine, NEAA (1%), HEPES (1 µM) and 50 µg/mL Penicillin streptomycin |
| NBM++ (conditioned medium) | 1x NBM, 1x B27, 2 mM L-Glutamine, and 25 µg/mL Gentamicin |

Primary rat embryo neurons were obtained from a rat Hippocampus on embryonic day 18 (E18). They were a gift from A. Püschel. All animal protocols were performed in accordance with the guidelines of the North Rhine-Westphalia State Environment Agency (Landesamt für Natur, Umwelt und Verbraucherschutz (LANUV)). Rats were maintained at the animal facility of the Institute for Integrative Cell Biology and Physiology (University of Münster) under standard housing conditions with a 12 h light/dark cycle (lights on from 07:00 to 19:00 h) at constant temperature (23°C) and with ad libitum supply of food and water. Timed pregnant rats were set up in house. Pregnant rats were anesthetized by exposure to isoflurane followed by decapitation and primary cultures prepared from embryos at embryonic day 18 (E18). During dissection, neurons from all embryos (regardless of sex) were pooled.

The growth medium for neurons comprised of 50 mL Neurobasal medium, 1 mL B27, 500 μl 100x Glutamine and 125 μl Gentamicin. This conditioned medium was used as growth medium. Neurons were kept at 37°C and 5% CO_2_. To obtain polar neurons, Hippocampi were isolated from rat embryos (at day 18 of embryonic development) and collected in HBSS on ice. Hippocampi were incubated at 37°C for 8 minutes in 2 mL Trypsin (not re-suspended). They were washed five times in pre-warmed DMEM++ before being resuspended in 2 mL DMEM++ with a 1 mL pipette tip after aspirating trypsin. Neurons were then seeded on poly-ornithine coated dishes with NBM++ medium and incubated at 37°C with 5% CO_2_. For microscopy, around 50,000 HeLa cells were seeded on a 3 cm glass cover slip or in a 3 cm glass bottom dish (Ibidi™). HeLa cells were seeded at least 24 hours before transfection. 100,000 neurons were seeded on poly-ornithine coated 3 cm glass bottom dishes. Neurons were cultured in DMEM++ medium, which was replaced with conditioned media after 3 hours and placed in an incubator at 37 °C and 5% CO_2_. The day when neurons were seeded is termed as “Day in vitro zero” (DIV0). The first day following seeding is known as DIV1, and the days after that are known as DIV2, DIV3, and so on.

### Coverslip coating for neurons and transfection

The dishes were coated under a cell culture hood. A 1:100 dilution of poly-ornithine in PBS was used to coat 3 cm Ibidi dishes. Dishes were incubated for 1 hour at 37 °C. Lipofectamine 3000 was used to transiently transfect HeLa cells according to the manufacturer’s instructions. After transfection, the cells were cultured overnight in an incubator at 37 °C and 5% CO_2_. The following day, live HeLa cells were imaged using TIRFM. For neuronal transfection, the conditioned medium was carefully removed from the glass bottom dishes with neurons and stored at 37 °C. DNA was delivered to neurons by the calcium phosphate transfection procedure. After being washed, neurons were supplied with pre-collected conditioned medium and left for three days to mature. At DIV3, live neurons were imaged.

### Staining self-labeling tags for live cell imaging

The protein of interest (POI) was genetically fused to either the HaloTag or the Snaptag at the C-or the N-terminus of the Exo70 protein sequence resulting in Exo70-HaloTag (C-tagged) or HaloTag-Exo70 (N-tagged). The HaloTag was stained with a dye that was coupled to a derivative of 1-chlorohexane as a substrate. This is called the HaloTag ligand (HTL) and it forms a covalent bond with the HaloTag. A benzyl-guanine (BG) was coupled to a dye for staining the SnapTag substrate. The fluorescent dye forms a covalent thioether bond with the SnapTag via the benzyl-guanine. Janelia Fluor® (JF) dyes JF646-HTL and JF549-BG were used for staining the respective tags. In the case of dual-color experiments, one POI was fused to the HaloTag and the other POI to the SnapTag. Hence for dual color experiments labelling was performed by combining JF-646-HTL and JF-549-BG. Halo-tagging outperformed Snap tagging, resulting in brighter structures achieved with lower concentrations ^32^. In dual-color microscopy, different dye labelling concentrations were utilized due to the differing binding kinetics of the two tags.

Transfected cells were stained just before imaging. Sub-stoichiometric labelling was used for single molecular studies performed with the TIRF microscope. Dyes were diluted in imaging medium a final concentration of 500 picomolar (pM) of JF646-HTL. JF549-BG was used in a final concentration of 2 nano-Molar (nM). Cell dishes were washed in warm PBS once, then stained with respective tagging dyes for 30 minutes at 37°C and 5% CO_2_. Cells were then rinsed three times with pre-warmed PBS after which pre-warmed imaging medium was added. After staining, cells were allowed to rest in the incubator for at least 15 minutes before being imaged. To obtain higher labeling densities for confocal imaging 50 nM JF646-HTL and 50 nM JF549-BG were used to stain the cells. Cells were incubated for 30 minutes and later rinsed three times with PBS.

### Immunostaining

Cells were fixed with 4% paraformaldehyde (PFA) containing 15% sucrose for 10 minutes at room-temperature (RT). Following PFA fixation, samples were washed three times in PBS before being treated with 50 mM Ammonium Chloride (NH_4_Cl) to eliminate free aldehyde groups. Then, cells were permeabilized with 0.1% TritonX-100 in PBS for 10 min. After washing with PBS, unspecific antibody binding was minimized by blocking with 2% goat serum and 3% bovine serum albumin (BSA) (1 hour at room temperature in a dark, humid environment). Any residual blocking buffer was aspirated after 1 hour. Both HeLa cells and neurons were immune-stained with a primary antibody against endogenous Exo70 (12014-1-AP, Proteintech) at a dilution of 1: 50. The secondary antibody was conjugated with Alexa Fluor 555 (A-21422, Thermofischer) and used at a dilution of 1: 500. A negative control without the primary antibody was set up at the same time. The samples were kept in the dark for an overnight incubation at 4 °C. To remove unconjugated antibodies, cells were washed three times with PBS. For WESTERN, antibodies listed in Table 2 were used.

**Table 2:**
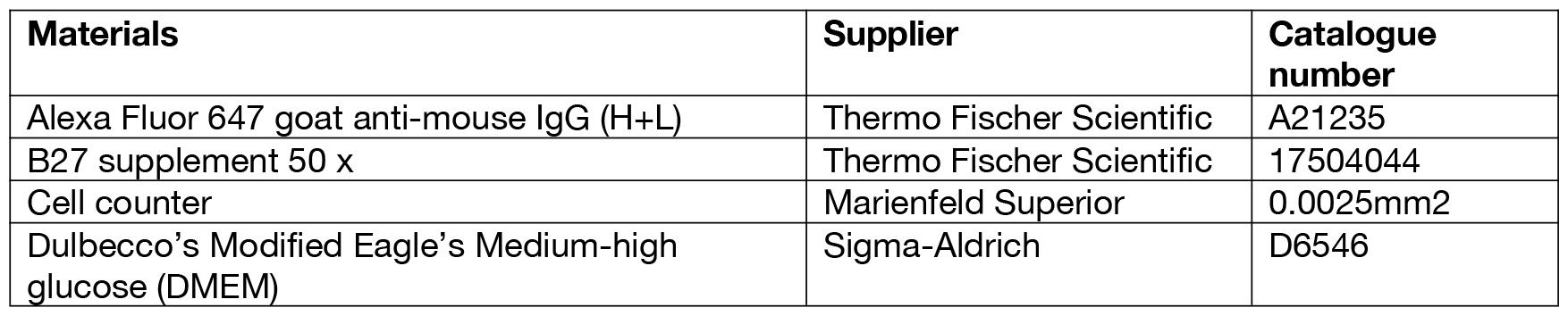
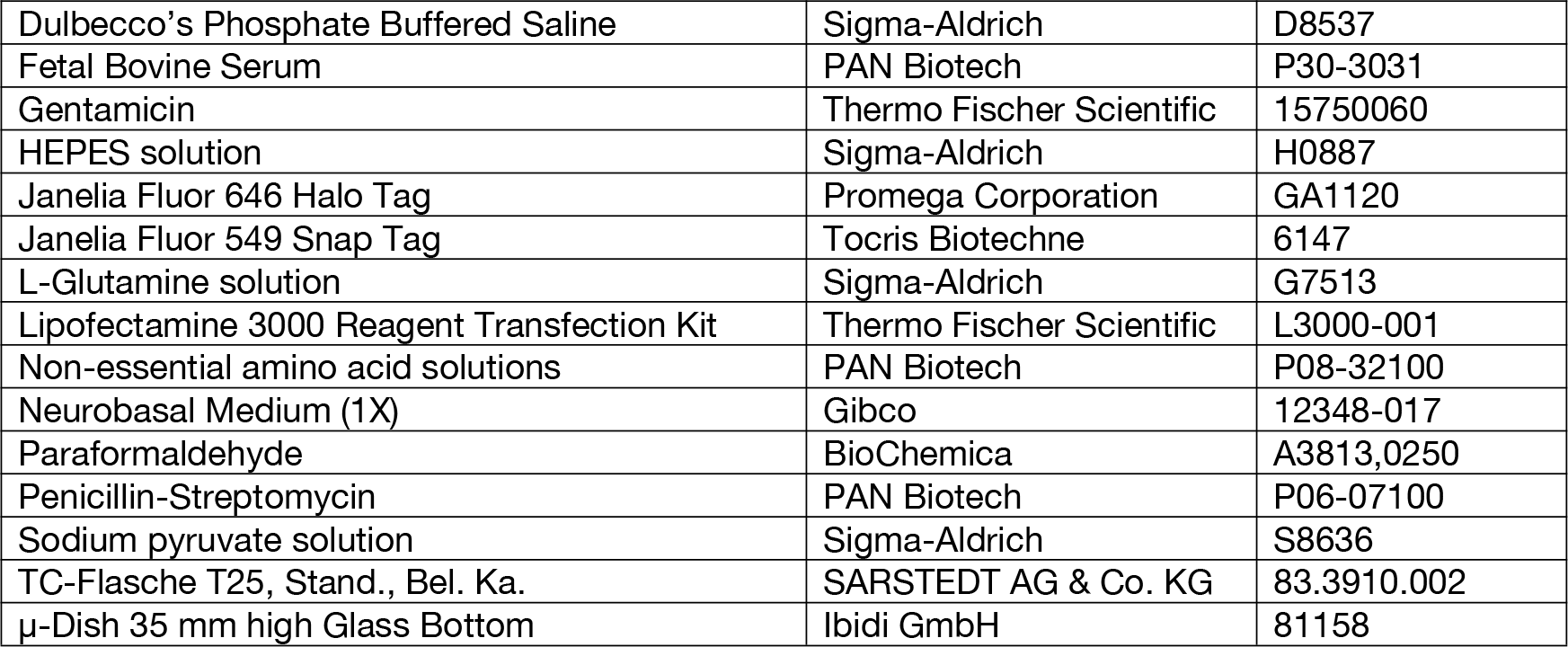
General Material

**Table 3:** List of antibodies for Immunofluorescence staining

| Antibody target | specification | Provider | Ordering number |
| --- | --- | --- | --- |
| Exo70 | Mouse monoclonal | Santa Cruz | sc-365825 |
| Sec 3 | Rabbit Polyclonal | Thermo Fischer Scientific | A303-363A |
| Alexa Fluor 488 | goat anti-rabbit | Thermo Fischer Scientific | A-11008 |
| Alexa Fluor 555 | goat anti-mouse | Thermo Fischer Scientific | A21147 |

### Cloning of sfGFP-Exo70 and Exo70-sfGFP

The N-terminus Exo70 (EXOC7) was fused to sfGFP to generate sfGFP-Exo70. SfGFP was amplified from psfGFP-C1 using the primers 5’-cgcaaatggc gcgtaggcgtg-3’ and 5’-CAGATCTGAG TCCGGCCGGA CTTGTACAGC TCGTCCATGC-3’ and inserted into pEGFP-C3-Exo70 (Martin et al., 2014) using NheI and BglII to replace EGFP. Exo70-sfGFP was constructed by fusing the C-terminus of Exo70 to sfGFP. Exo70 was amplified from pEGFP-C3-Exo70 using the primers 5’-CGGAATTCTG ATGATTCCCC CACAGGAGG-3’ and 5’-GCGTCGACAC GGCAGAGGTG TCGAAAAGGC-3’ and inserting the product into psfGFP-N1 using EcoRI and SalI. psfGFP-C1 was a gift from M. Davidson and G. Waldo (Addgene plasmid #54579; http://n2t.net/addgene:54579; RRID: Addgene_54579) and pEGFP-C3-Exo70 from C. Der (Addgene plasmid #53761; http://n2t.net/addgene:53761; RRID: Addgene_53761).

### Confocal microscopy

For confocal and fluorescence lifetime imaging, the commercial inverted microscope, Leica TCS SP8 FLIM system, was used. The microscope was equipped with a unique Acoustic Optical Deflector (AOD), two photomultiplier tubes (PMT) and two highly sensitive hybrid GAsP (HyD) detectors. A 63x water objective (NA 1.2) was used for imaging. The configuration includes a pulsed white light laser (WLL) that offers up to eight fully tunable excitation lines between 470 and 670 nm simultaneously. The WLL was also used for time correlated single photon counting, which was controlled by the Picoquant™ software. The microscope was equipped with a temperature control, gas regulation, and a humidifier. An image format of 1024x1024 pixels was used with a scan speed of 700 Hz to obtain images with high signal to noise ratio. The immunostained endogenous Exo70 in cells was visualized by excitation with the white light laser tuned at a wavelength of 555 nm, and a z-stack of images was acquired. Stacks were then processed in FIJI to create an image that represented an averaged intensity projection of all the images in the stack.

### TIRF microscopy

An inverted TIRF microscope (Xplore, Olympus) was used for SPT studies. The microscope is equipped with a TIRF objective (100x oil, NA of 1.49), an Optosplit IV for simultaneous recording of four channels and four diode lasers (wavelengths of 405 nm, 488 nm, 560 nm, and 640 nm) connected by an optical fiber to four independently controlled TIRF modules. To reduce wide-field illumination, a collimated beam of light was generated by focusing into the back focal plane of the objective. The output intensities of the diode laser in the system were controlled on a microsecond timescale by an acoustical optical tunable filer (AOTF). To image single molecules, we used a highly sensitive sCMOS camera (ORCA Fusion–BT, Hamamatsu) at a pixel size of 65 nm. For controlling imaging conditions, the CellSens software was used to set up automatic time series of 5,000 frames with a constant time interval (58.8 Hz). In this study, the image format for a single channel was set to 500x500 pixels. Each cell in a sample was recorded for a total of 85 seconds, and each technical replicate consisted of 10 cells in total.

### Single particle tracking and data processing

For tracking analysis, the raw images were processed in FIJI using the Trackmate plugin to extract track information for the analysis ^26^. This program takes the simplest approach possible by detecting each particle’s nearest neighbors in a circular region in successive frames. Before utilizing the plugin, it was double-checked that FIJI’s image properties were correct. Spots were detected using the “LoG detector” filter. In combination with the median filter and subpixel localisation, the threshold was modified according to the signal intensity. The data was visualized using the “HyperStack Displayer” and uniform color. The tracking was done using the simple LAP tracker (linking conditions: maximum distance, gap-closing conditions, maximum distance). The generated tracks were further filtered using a “Number of spots in tracks” filter. This filter was used to make sure the tracks were long to reduce statistical errors and discard tracks produced by blinking occurrences. The resulting tracks were saved as a “.XML” file, which could then be further analyzed in Matlab using the @msdanalyser to get ensemble MSD curves, diffusion coefficient distributions, and alpha coefficient distributions ^33^. Generally, ensemble MSD curves were plotted after weighted averaging over all particles. Diffusion coefficients were extracted from the MSD curves using power law and fitted with a linear function. The first 25% of the trajectory were fitted. The threshold for the quality of the fit were R2 coefficients > 0.8. The fits gave us the values of a and 𝒟 for individual tracks according to the models described by Qian ^34^. For statistical analysis, Origin™ software was used for analysis of and 𝒟.

### TCSPC

Lifetime imaging was performed with the confocal LSM to capture lifetime decays in each pixel as time-correlated single photon counting (TCSPC). Therefore, a PicoQuant HydraHarp 400 TCSPC unit in conjunction with a Leica SP8 confocal microscope was employed. The Hydra Harp 400 had a count rate of up to 12.5 million counts per second per channel and a temporal resolution of 1 picosecond (ps). A pulsed WLL (set to λ=488 *nm*) with a 40 MHz repetition time rate was used as the excitation source in combination with a HyD detector. The simulated instrumental response function (IRF) had a width of 150 ps. An emission filter BP525/50 nm was used. Fluorescence was recorded between 500 − 540 *nm* with 16 *ps* time interval. The acquisition was performed until at least 1,000 photons in the brightest pixel were reached. PicoQuant’s TCSPC electronics in the confocal microscope was controlled by SymPhoTime 64, a customized data acquisition software. This software was also used for TCSPC lifetime decay fitting with advanced error correction and included mono- and bi-exponential fitting of fluorescence decay curves (after subtracting the IRF) from the region of interests. The TCSPC curves were analyzed using the SymPhoTime 64 software. The “n-Exponential Reconvolution Fit” model was selected as the final fit for the TCSPC histogram considering the simulated IRF. For the calculation of the average lifetime τ, we used the intensity weighted average lifetime “τ_Av_Int_”.

### Lattice light-sheet microscopy

Lattice light-sheet microscopy was performed on a home-built clone of the original design by the Eric Betzig group (Chen et al, 2014). Samples were prepared as following: ∼ 140,000 neurons (DIV0) were seeded on glass coverslips (5 mm, Art. No. 11888372, Thermo Scientific). Transfection of cells were conducted with the Calcium phosphate transfection method using 1.5 μg DNA. After 1 h, neurons were rinsed with pre-warmed OpitMEM solution. Cells were cultured until DIV3 and then fixed using 4 % paraformaldehyde solution. The glass coverslips with the fixed cells were inserted into a custom build sample holder that was attached on top of a sample piezo. This ensures that the sample is inserted at the correct position between the excitation and detection objectives inside the sample bath containing PBS pH 7.4 at room temperature. For image acquisition, an image stack was acquired in sample scan mode by scanning the sample through a fixed light sheet with a step size of 350 nm, which is equivalent to a ∼ 190 nm slicing with respect to the *z*-axis considering the sample scan angle of 32.8°. A dithered square lattice pattern generated by multiple Bessel beams using an inner and outer numerical aperture of the excitation objective of 0.48 and 0.55 was used during the experiments. Excitation was done by a 488 nm laser (2RU-VFL-P-300-488-B1R; MPB Communications Inc., Pointe-Claire, Canada). Emission was collected by a water dipping objective (<u>CFI Apo LWD 25XW</u>, NA 1.1, Nikon) and finally recorded by a sCMOS camera (ORCA-Fusion, Hamamatsu, Japan) with a final pixel size of 103.5 nm and an exposure time of 50 ms. Raw datasets were further processed with an open-source LLSM post-processing utility called LLSpy (https://github.com/tlambert03/LLSpy) for de-skewing and deconvolution. Deconvolution was performed by using an experimental point spread function recorded from 100 nm sized FluoSpheres™ (Art. No. F8803, ThermoFisher Scientific) and is based on the Richardson–Lucy algorithm using 10 iterations.

### Molecular modeling procedure

3D homology models for Exo70-GFP-fusion constructs were generated as following: Initially, partial Exo70 models were constructed by the Swiss-Model web-server whereas the precise amino acid sequences of the Exo70 plasmids served as input. The Exo70 homology model was built alongside GFP (4KW4.pdb), and flexible linking sequences (*de novo* construction by YASARA Structure ver.21.12.19) were also included in the homology model. The resulting models represented structural models of the Exo70 N- or C-terminal tagged constructs encoded by the plasmids used here (depicted in Figure 5 a,b). Exo70 contains glycine linkers at the C-terminus, allowing for some flexibility and enabling it to stretch to adapt to a required shape. However, the movements of Exo70 tagged at the N-terminus were somewhat restricted. Both, the N- or C-terminal GFP tagged Exo70 constructs, were extensively energy minimized using YASARA Structure (ver.21.12.19). The two models GFP fused Exo70 models were then used to substitute Exo70 in the yeast homologue of the exocyst complex from a recently published the Cryo-EM-structure (5YFP.pdb). During this interactive docking process, the glycine-linkers were kept flexible to allow for efficient incorporation of the GFP fused Exo70 models. However, *in silico* docking shows that Exo70 tagged at the C-terminus can effectively be incorporated into the complex, whereas the GFP in Exo70 tagged at the N-terminus interferes with complex formation of *in silico (*Figure 4D) suggesting compromised complex formation in a native form as captured in the cryo-EM structure.

## 3 RESULTS AND DISCUSSION

### N-terminally tagged Exo70 induces filopodia formation in unpolarized cells and neurons

We first asked, where endogenous Exo70 localizes in cells. As immunostaining showed, Exo70 is cytoplasmic and at the cell membrane in unpolarized HeLa cells and in neurons. Next, we compared the localization of N- and C-terminally tagged Exo70 and found distinct distributions and functions (Figure S1). While Exo70-HaloTag showed a homogenous cytosolic distribution, HaloTag-Exo70 accumulated in spots at the plasma membrane in addition to its cytoplasmic localization. HaloTag-Exo70 also induced the formation of filopodia in Hela cells as described earlier for GFP-Exo70 ^11,14^. Also, hippocampal neurons derived from rat embryos at day 18 and transfected with Exo70-HaloTag and HaloTag-Exo70, respectively, show clearly different localization of tagged Exo70. N-terminally tagged Exo70 was found in spots in neuronal compartments in DIV1, DIV2 and DIV3, and extra membrane extensions in the soma and throughout the axon were observed already in DIV1 (Figure 1A). These filopodia structures became more apparent at DIV2 and DIV3 (Figure 1B, C). In contrast, cellular morphology was not affected by overexpression of Exo70-HaloTag, which was homogenously distributed in neurons (Figure 1D-F). To visualize the effects of N-terminal tagging of Exo70 with more detail, we imaged sfGFP-Exo70 in DIV3 neurons by lattice light sheet microscopy. sfGFP-Exo70 was found in the axon cytoplasm (Figure 1G-I) but also at the plasma membrane (Figure 1G-I, yellow arrowheads). The growth cone (GC) displays a high concentration of sfGFP-Exo70. Moreover, multiple membrane protrusions were induced in the growth cone and in the axon (Figure 1G-I, red arrowheads). Also, cells with manifold cell extensions were found that exhibited no prominent axon (yet) (Figure 1H). These observations confirm a role of sfGFP-Exo70 in filopodia formation, likely through interaction with the Arp2/3 complex, as suggested by previous studies 15,16. Exo70-sfGFP on the other hand, had no apparent effect on normal neuron morphology. The protein was mainly found in the cytoplasm (Figure 1J-L) high-resolution images clearly demonstrate the distinct effect of sfGFP-Exo70 shaping neuronal, in particular axon morphology. sfGFP-Exo70 appears to be hyperactive in terms of promoting membrane extension.

**Figure 1:**
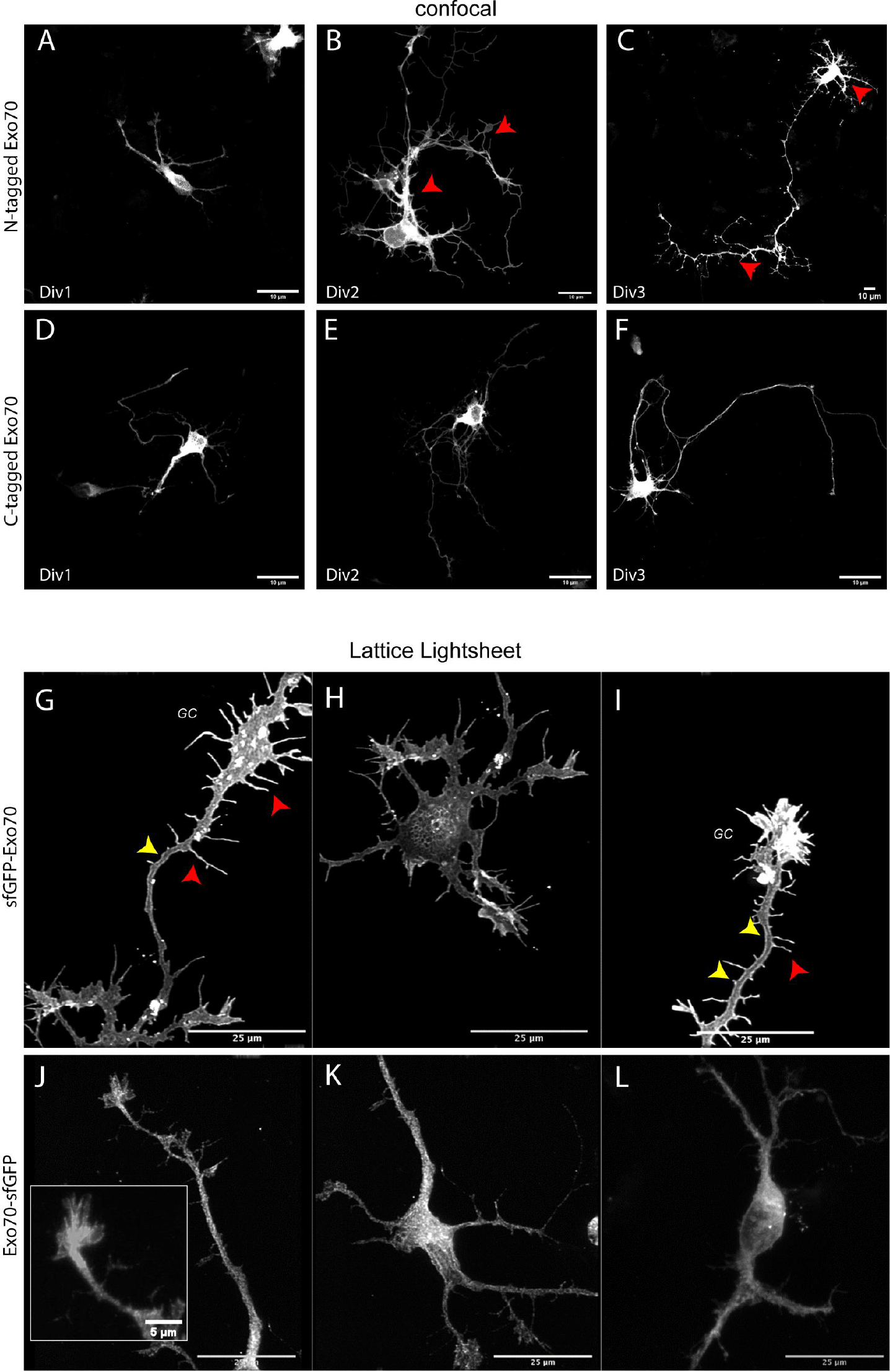
Subcellular localization and effects of Exo70-sfGFP and sfGFP-Exo70 in rat hippocampal neurons. (A-C) Confocal images of HaloTag-Exo70 in the cytosol of the soma, in dendrites and axons of neurons at DIV1, DIV2 and DIV3. z-stacks were recorded, and the intensity averaged (FIJI). From DIV2 onwards, red arrow heads indicate spine formation. (D-F) Exo70-Halo distribution neurons at DIV1, DIV2 and DIV3. (G-I) Lattice light sheet micrographs of neurons expressing sfGFP-Exo70 at DIV3. (J-L) Growth cone (framed) and axon of a neuron transfected with Exo70-sfGFP. Scale bars: 10 μm (A-F), 25 μm (G-I), 5 μm (inset (J).

### Exo70-sfGFP’s decreased fluorescence lifetime suggests its involvement in complex formation

Exo70/exocyst complex tethers vesicles to the PM. Therefore, the nano-environment of Exo70 is expected to provide electro-dynamic interactions which should reduce the fluorescence lifetime (τ) of an attached fluorophore. In contrast, Exo70 interacting with Arp2/3 during membrane extension encounters a different nano-environment. To test, whether N- and C-terminally fused sfGFP would exhibit differences in their fluorescence lifetime τ, we performed fluorescence lifetime imaging microscopy (FLIM) using time correlated single photon counting (TCSPC). In the FLIM image, Exo70-sfGFP shows a homogenous distribution, while sfGFP-Exo0 was less homogenously distributed. Moreover, strong filipodia formation was observed (Figure 2A). The color code already indicates difference in τ between Exo70-sfGFP and sfGFP-Exo70. As demonstrated with an exemplary lifetime decay curve, the decay curve of sfGFP−Exo70 was shifted towards longer lifetimes compared to Exo70-sfGFP (Figure 2B). Bi-exponential fitting of fluorescence decay curves (after subtracting the IRF) resulted in two lifetimes, that are plotted for each construct comparing (Figure 2C). Average τ _AV-INT_ and τ _AV-AMP_ values were compared. sfGFP-Exo70 displayed a 0.09 ns shorter lifetime τ _AV-AMP_ than sfGFP-Exo70 (*p*<0.001). The average lifetime τ_AV-INT_ of sfGFP in the two constructs differed by 0.11 ns (*p*<0.001) (Figure 2C). The lower τ_1_ and τ_AV int_ of Exo70-sfGFP suggests that it is exposed to a molecular environment that provides more electro-dynamic interactions^17^, such as an Exo70-sfGFP/exocyst complex assembly that tethers vesicles to the plasma membrane. In contrast, the longer lifetime of sfGFP-Exo70 indicates that less Exo70 is assembled.

**Figure 2:**
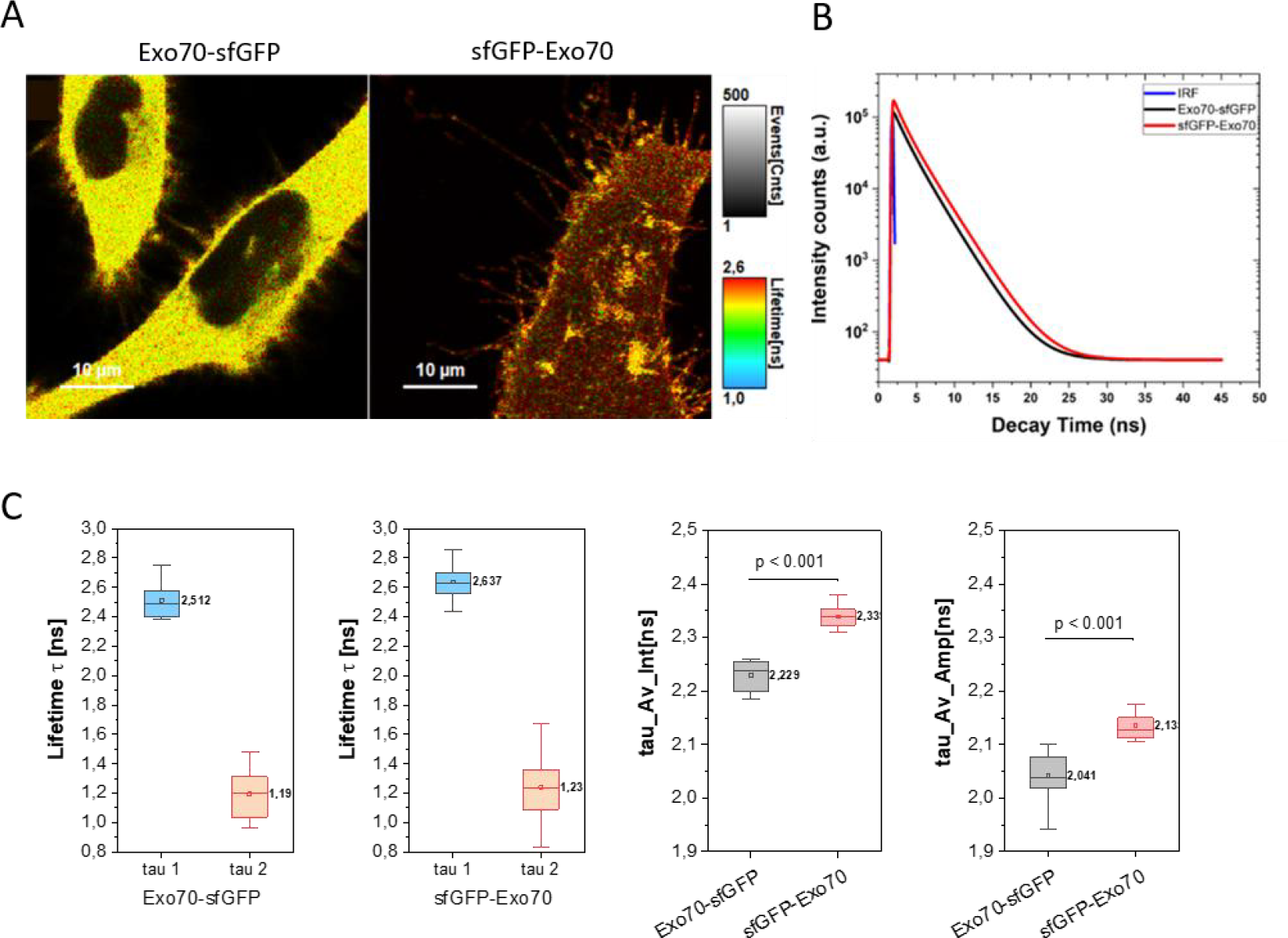
sfGFP-Exo70 and Exo70-sfGFP exhibit different nano-environment. (A-B) False colored fluorescent lifetime maps for Exo70-sfGFP (A) and sfGFP-Exo70 (B) in HeLa cells. (C) An exemplary TCSPC curve is displayed as semi-logarithmic plot of sfGFP fluorescence lifetime decay; comparison of sfGFP fused C-terminal (black line) and N-terminal (red line) to Exo70. Blue line: instrumental response function (IRF). (C) Lifetimes of sfGFP-Exo70 and Exo70-sfGFP. Statistics: Box-and Whiskers blots: 25-75%, median as straight line, numbers indicate mean values, ANOVA one way using Tukey test. Exo70-sfGFP n=11, sfGFP-Exo70 n=15 cells. Scale bars: 10 μm (A, B).

### Single particle tracking shows higher mobility of N-terminal tagged Exo70 in HeLa cells

Next, we tested whether we could distinguish different subpopulations of Exo70 by monitoring its mobility at the plasma membrane. Therefore, single particle tracking (SPT) with HaloTagged Exo70 versions was performed using TIRF illumination. The HaloTag is commonly used in single particle tracking and localization microscopy ^13,18^. HeLa cells were transfected with Exo70-HaloTag and HaloTag-Exo70, respectively, and an empty vector as a control cell line. After sub-stoichiometric labeling with the fluorescent Halo-Tag-Ligand JF646-HTL (500 pM), images were captured in a TIRF microscope equipped with a highly sensitive sCMOS camera. In the epi-fluorescent excitation mode, Exo70-HaloTag/JF646-HTL showed homogenous distribution (Figure 3A) as found before (Figure S1, Figure 2A). Also, the HaloTag-Exo70 induced membrane extrusions were clearly visible (Figure 3B). In the TIRF mode, single particles were discernable in both the cell body and in distinct filopodia.

**Figure 3:**
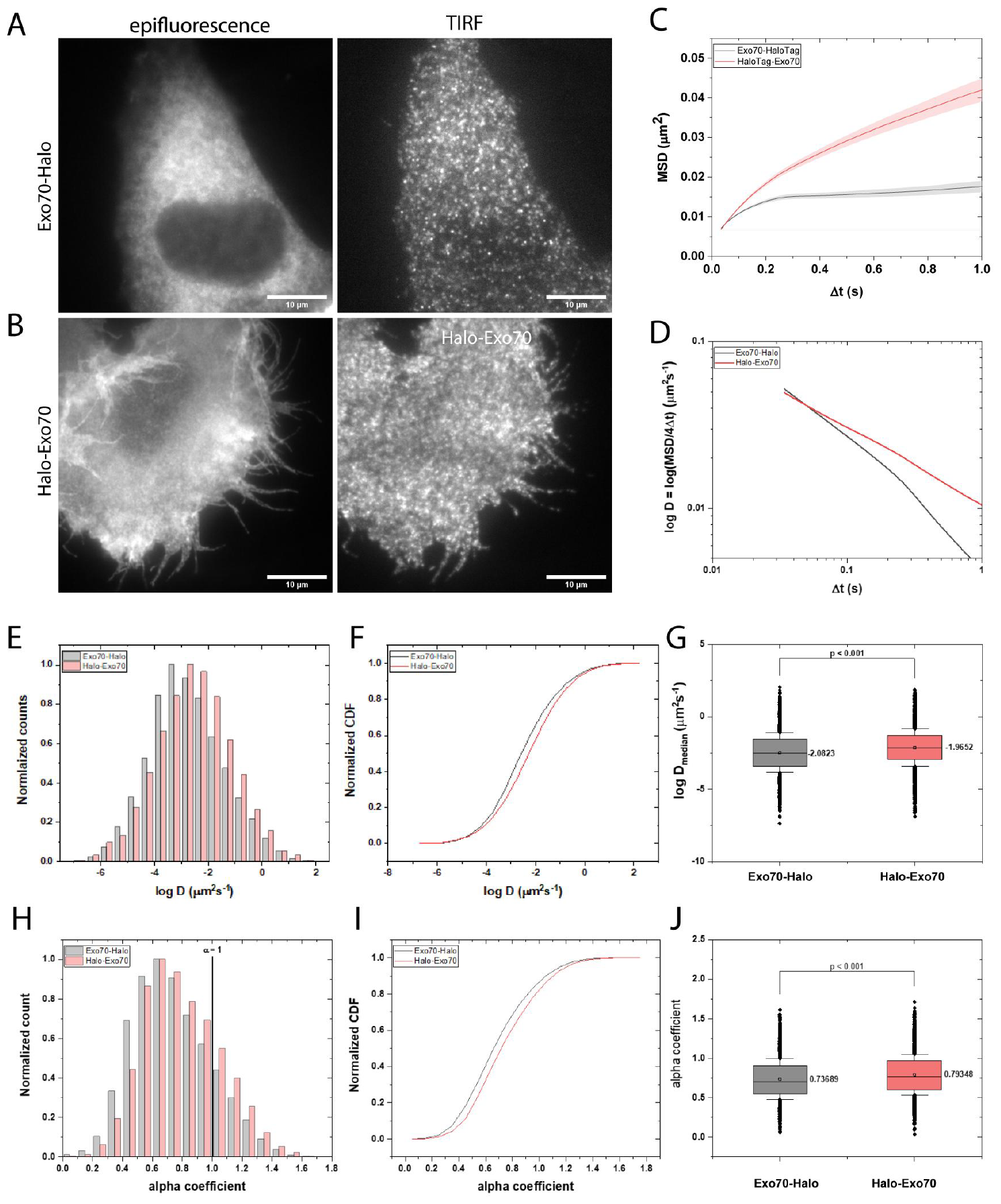
N-terminally tagged Exo70 shows higher mobility than C-terminally tagged Exo70 in HeLa cells. (A) Images of Exo70-HaloTag (JF646-HTL) transfected HeLa cells taken in epifluorescent and TIRF excitation mode. (B) Images of HaloTag-Exo70 (JF646-HTL) transfected cells taken in epi-fluorescent and TIRF excitation mode, respectively. (C) Mean square displacement of Exo70-HaloTag and HaloTag-Exo70 (JF646-HTL). (D) Changes of the diffusion coefficient (log D) with increasing time intervals Δt. (E) Distribution of the diffusion coefficients (as log D). (F) Cumulative density function (CDF) of diffusion coefficients. (G) Box and Whiskers (25th and 75th percentiles) inclusive outliners showing the median diffusion coefficients (as log D), with standard error of the man (S.E.M.). (H) Normalized histogram of the alpha coefficient distribution for Exo70-HaloTag (grey bars) and HaloTag-Exo70 (red bars). A straight line is plotted at = 1. (I) CDF of Exo70-HaloTag (black line) and CDF of HaloTag-Exo70 (red line). (J) Comparison of alpha coefficients of Exo70-HaloTag diffusion and that of HaloTag-Exo70 diffusion. The p values were calculated using Kruskal-Wallis Anova and the design of the box are through Tukey test. N=3 (all data e-i). Scale bars: 10 μm (a,b).

The Exo70 particles were localized and tracked using the FIJI plugin TrackMate ^19^. The tracks from three similar SPT experiments were pooled to obtain ensemble time averaged mean square displacement (MSD) curves as shown in Figure 3 (C). The ensemble-MSD curves show that Exo70-HaloTag (black curve; 13,243 tracks) exhibited confined motion while HaloTag-Exo70 (red curve; 13,085 tracks) exhibited anomalous behavior. When the diffusion coefficients are plotted in dependence on averaged delay time (Δt) on a logarithmic scale as shown in Figure 3 (D), the curves for Exo70-HaloTag and HaloTag-Exo70 are not parallel to the x-axis, confirming non-Brownian motion ^20^. Moreover, the kink in the black and red diffusion curves to a steeper negative slope indicates that Exo70-HaloTag and HaloTag-Exo70 undergo transition from anomalous to confined motion, suggesting two different states of Exo70 within a specific cellular environment. The probability density functions of the distribution of diffusion coefficients for Exo70-HaloTag and Halo-Tag-Exo70 shows a shift of Exo70-HaloTag particles towards slower diffusion (Figure 3E), which also visible in the cumulative density function (CDF) plot (Figure 3F). The CDF curves for Exo70−Halo (black curves) show that the diffusion coefficients were lower than for HaloTag-Exo70 (red curves). Finally, the diffusion coefficients for all tracks of Exo70-HaloTag and HaloTag-Exo70 were compared (Figure 3G). For HaloTag-Exo70, the diffusion coefficient was *D* = 0.011 ± 0.015 μm^2^/s (median ± s.e.m) and for Exo70-HaloTag *D* = 0.008 ± 0.017 μm^2^/s (median ± s.e.m). These low diffusion coefficients indicate that the majority of Exo70 on the plasma membrane is quasi-immobile as expected for example for Exo70 in the PM-tethered exocyst/vesicle complex. The diffusion coefficient for Exo70-HaloTag at the plasma membrane was ∼24% lower than that of HaloTag-Exo70, implying that more Exo70-HaloTag is associated to the exocyst/vesicle tethering complex than HaloTag-Exo70.

In order to quantify subpopulations of Exo70, we determined alpha (α) coefficients, which describe the diffusion mode. An alpha coefficient of α = 1 indicates Brownian diffusion, α < 1 anomalous diffusion and α > 1 directed motion. The mean alpha coefficient α_Exo70-HaloTag_ = 0.74 ± 0.003 was slightly smaller than α_HaloTag-Exo70_= 0.79 ± 0.003 (Figure 3H-J). This in line with more restricted diffusion of Exo70-HaloTag. 76% of Exo70-HaloTag molecules and 70% HaloTag-Exo70 molecules displayed anomalous or confined motion, supporting the model that less HaloTag-Exo70 is engaged in interaction with exocyst/vesicle tethering.

### 3D Homology modeling of Exo70 tagged to GFP

The SPT and FLIM results suggest that Exo70-HaloTag is preferred to HaloTag-Exo70 in the formation of exocyst complexes. To address, whether the tag at the N-terminus might interfere with Exo70/exocyst assembly, 3D homology models were created for GFP-Exo70 and Exo70-GFP in the yeast exocyst complex (5YFP). We used the Swiss pdb software and the amino acid sequences of the Exo70 plasmids. First, the Exo70 homology model was built with GFP (4KW4) including linker sequences, whereby GFP (red beta sheets) was fused to Exo70 at the C-terminus (Figure 4A) and the N-terminus (Figure 4B), respectively. The alpha-helical structure is represented by the blue barrels (total 19), while the linkers are shown by the green lines. Exo70 contains glycine linkers at the C-terminus, making it more mobile in its movements and enabling it to stretch to adapt to a required shape. However, the movements of Exo70 tagged at the N-terminus were restricted. GFP fused to Exo70 homology model was then used to substitute Exo70 in the yeast homologue of the exocyst complex based on the Cryo-EM-structure (5YFP). The GFP-fused Exo70 is illustrated in red, while the other seven exocyst complex subunits from yeast are shown in blue in Figure 4. As the modelling shows, Exo70 tagged at the C-terminus can be effectively incorporated into the complex, (Figure 4C),while GFP tagging of the N-terminus interfered with Exo70/Exocyst complex formation (Figure 4D) as predicted earlier ^6^.

**Figure 4:**
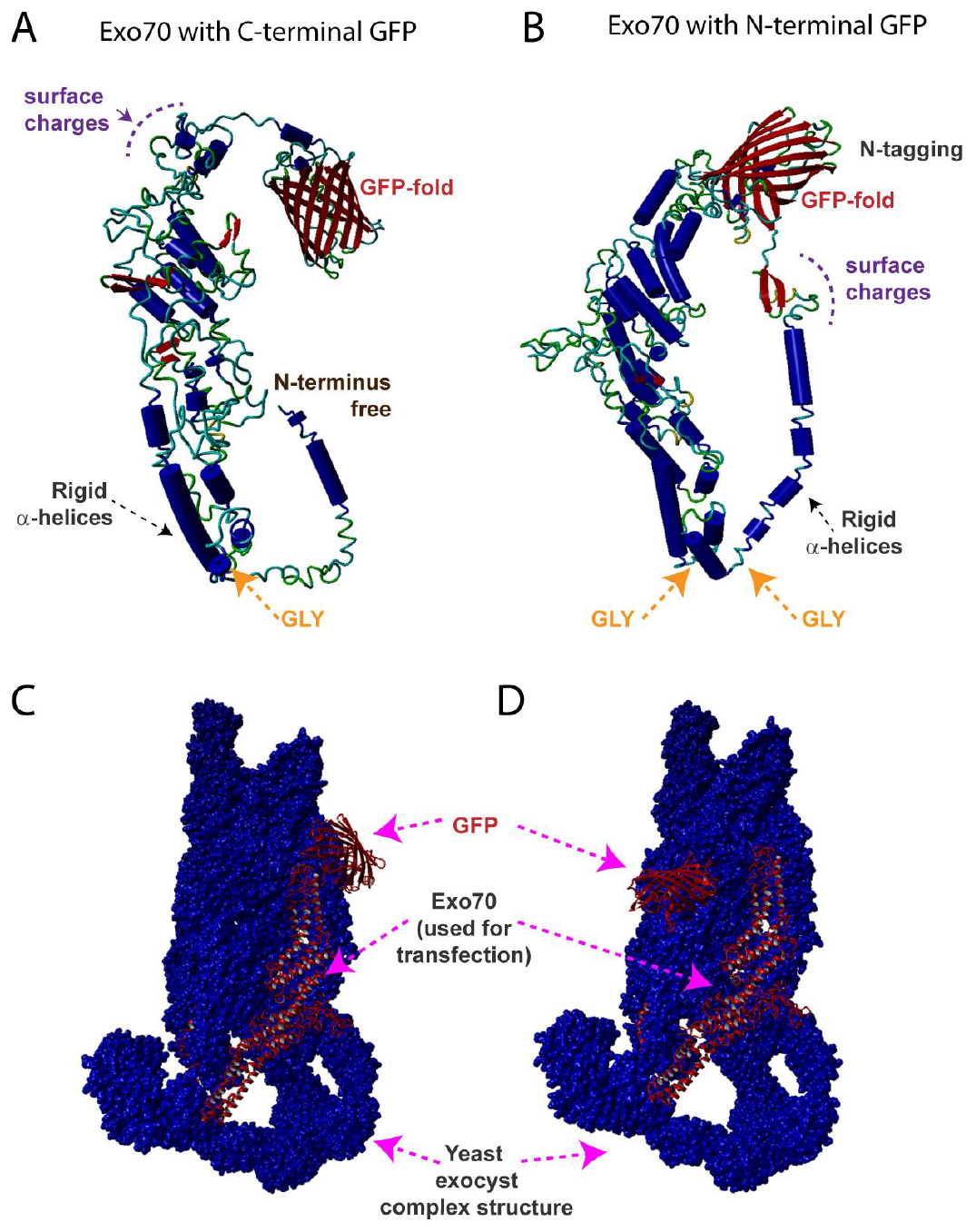
Homology models of Exo70-GFP and GFP-Exo70 and their integration into an exocyst complex. 3D model of Exo70 when GFP is tagged to the (A) C-terminus and (B) N-terminus. The red sheets in the barrel represent GFP. The blue cylinders indicate alpha helical coils, and the green lines indicate the linkers. (C) Exo70-GFP (red color) incorporated in the structure of the exocyst complex from yeast shown in blue color. (D) GFP-Exo70 interferes with assembly into the exocyst complex. The entire complex was created by minimizing the energy levels.

### Exo70-HaloTag interacts with exocyst subunit Sec3 indicating vesicle tethering

Finally, to test for Exo70-HaloTag assembly into the exocyst complex, the interaction with subunit Sec3 was checked. Exo70 and Sec3 interact with phosphatidylinositol 4,5-bisphosphate (PI(4,5)P2) in the plasma membrane and with vesicles at the same time ^21-23^. Colocalization of Exo70 and Sec3 therefore indicates exocyst/vesicle tethering to the plasma membrane. Immunostaining shows the co-localization of endogenous Sec3 (green) and Exo70 (red) at the plasma membrane Figure 5 (A-E). A detailed zoom in view (Figure 5D) displays various white punctae indicating colocalization of Exo70 and Sec3 (some of them are marked with pink arrows). The Pearson’s correlation coefficient calculated from the scatter plot in Figure 5 (E) was *r* = 0.497. This is in line with previous findings, where it was estimated that roughly 65% of subunits are assembled into the full exocyst complex ^21^.

**Figure 5:**
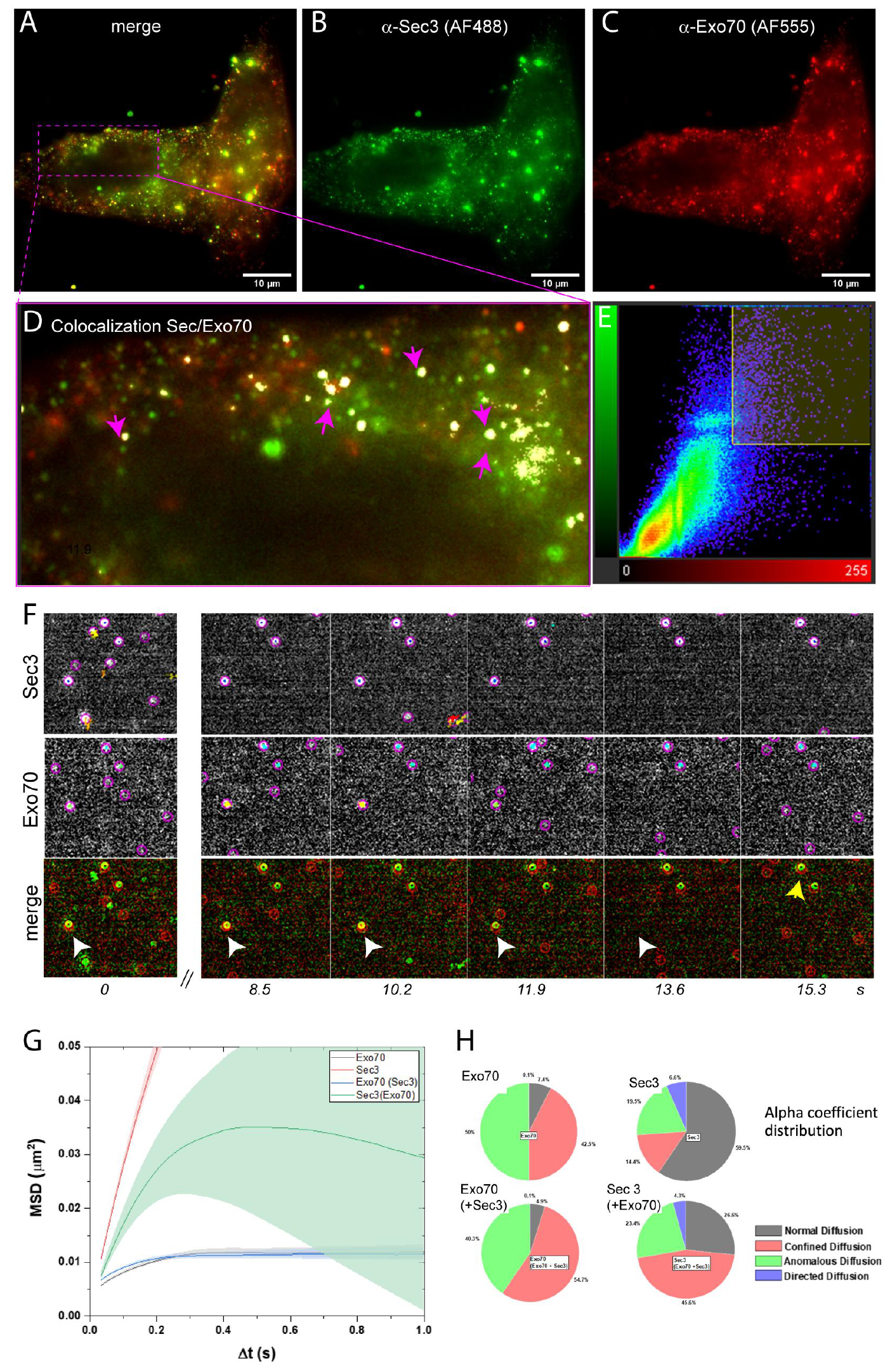
Colocalization and -migration of Exo70 and Sec3 in HeLa cells at the plasma membrane. (A-D) TIRF images of immunostained endogenous Exo70 and Sec 3. (A) Merged channels of Sec3 and Exo70. (B) Sec3 (green, 2^nd^ antibody labeled with AF488), (C) Exo70 (red, 2^nd^ antibody labeled with AF555). (D) Zoom in view of framed magenta box in (A). After threshold adjustment for both channels, co-localization analysis was performed in Imaris™. The white spots indicate Sec3 and Exo70 co-localization. (E) Scatterplot to determine the Pearson’s coefficient. Scale bars: 10 μm (a-c), 5 μm (d). (F) Time course showing co-localization of Sec3 and Exo70 particles for several seconds (white arrowhead, frame rate: 17ms). (G) Ensemble average MSD of Exo70 and Sec3 alone or in presence of the Sec3 and Exo70, respectively, in HeLa cells. (H) Quantification of Exo70 and Sec3 diffusion. Diffusion behavior was classified via α-coefficients, depicted in pie charts. Exo70: Exo70-HaloTag diffusion, Sec3: Sec3-Snaptag diffusion; Exo70(+Sec3): Exo70-HaloTag diffusion when Sec3 was overexpressed; Sec3(+Exo70), Sec3-SnapTag diffusion when Exo70 was overexpressed.

To check for interaction, dual color single molecule localization and tracking of Exo70-HaloTag and Sec3-SnapTag was conducted. The SnapTag was fused to the C-terminus of Sec3, since the N-terminus of Sec3 is crucial for the formation of the exocyst holo-complex ^6^.

HeLa cells were co-transfected with Exo70-HaloTag and Sec3-SnapTag and labelled with JF646-HTL (500 pM) and JF549-BG (2 nM), the substrate for the SnapTag. Dual color SPT with TIR excitation was applied to record the dynamics of Exo70 and Sec3 molecules at the plasma membrane until a maximal penetration depth of 150 nm. Single transfections were used as a control. Images were recorded with a frame rate of 58.8 Hz for a total of 85 seconds. An Optosplit IV device allowed for parallel recording of Sec3 and Exo70 in two channels.

Sec3 and Exo70 colocalization and comigration was observed in several cases (Figure 5F, bottom panel). Sec3 (green) and Exo70 (red), marked with a white arrow, colocalized from the start until 11.9 sec, then they disappeared together. In another case, Sec3 and Exo70, as marked with a yellow arrow, stayed during the entire 15.3 s recording time.

Finally, the impact of Sec3 on Exo70 diffusion and of Exo70 on Sec3 diffusion was quantified. The particles were tracked using FIJI plugin Trackmate and the resultant trajectories were sub-classified using another FIJI plugin Trajclassifier ^24^. MSD vs Δt diagrams show time ensemble average MSD for the different conditions (Figure 5G). The confined motion of Exo70 was only marginally affected by Sec3 overexpression indicating that Exo70 was already involved in exocyst/vesicle formation. On the other hand, Sec3, which exhibited mostly Brownian motion as alpha values indicate (Figure 5H), was significantly slowed down, when Exo70 was overexpressed. When Exo70 and Sec3 were co-expressed, Sec3’s confined population increased dramatically from 14.4% to 45%, suggesting that Sec3 requires Exo70 to be integrated into the exocyst complex. In consequence of this interaction, the fraction of anomalous diffusing Sec3 particles increased.

The dynamics of Sec3 is in line, which was observed earlier ^21^. The different diffusion modes of Sec3 and Exo70 suggest that only a fraction of Sec3 is assembled with Exo70/exocyst complex, while apparently the majority of Exo70 is found in the exocyst. Therefore, rather Exo70 serve as a landmark for exocytosis, and not Sec3 as suggested before ^4,22,25^.

## 4 CONCLUSION

Exo70 has a dual function in membrane expansion and as key subunit of the exocyst complex, which tethers exocytotic vesicles to the plasma membrane. In this study, we showed that the respective N- or C-terminal positioning of a tag (sfGFP or HaloTag) affects the subcellular localization of Exo70, its spatiotemporal dynamics and its function. This dissection was possible by high resolution imaging of Exo70. Exo70 is known to interact with PI(4,5)P2 at the plasma membrane via surface residues near its C-terminus, while the N-terminus is involved in regulatory interactions. Exo70 labeled at the N-terminus was present at the plasma membrane, but the data collected show a bias of N-terminal labeled Exo70 against exocyst complex formation (higher lifetime of a fluorescent tag, higher mobility). As 3D homology modeling revealed, that the formation of the exocyst complex is compromised with N-terminal tagged Exo70. Also, N-terminal labeled Exo70 strongly promoted filipodia formation in nonpolar (HeLa) and polar cells (neurons), which suggests interaction with Arp2/3. C-terminally labelled Exo70 showed restricted movement and reduced lifetime of an attached fluorophore, as expected for assembly into a plasma membrane-bound exocyst. In summary, the spatio-temporal organisation of Exo70 is influenced by the position of a C- or N-terminal tag and that this favours different interactions and functions of Exo70: Membrane expansion through exocyst formation or filipodia growth.

## 5 SUPPORTING MATERIAL

This part includes one figure.

## 6 AUTHORS CONTRIBUTIONS

HG and KB designed the experiments, HG conducted the experiments, GS and HG did the modeling, RK built the Lattice Light Sheet microscope, MH helped with Lattice Sheet experiments, HG and KB interpreted data and wrote the manuscript.

## 7 ACKNOWLEDGEMENTS

We thank Patrick Duwe and Frank Schmelter for technical assistance, Andreas Püschel and Priyadarshini Ravindran for valuable discussion and providing primary cultures of hippocampal neurons. The work was supported by the German Research Fundation (CRC) through a grant to H.G. (CRC 1348, # 386797833). Karin Busch and Andreas Püschel are members of the CiM (Cells in Motion Interfactulty Centre) at the University of Münster.

## 9 DATA

The datasets used and analyzed during the current study are available from the corresponding author on reasonable request.

